# Inositol-Requiring Enzyme 1 pathway and autophagy drive sequential response of endothelial cells to febrile range hyperthermia

**DOI:** 10.1101/2024.11.22.624904

**Authors:** Julie Vorbe, Florence Massey, Corinne Rocher, Océane Morales, Nihal Brikci, Marie Le Borgne, Giuseppina Caligiuri, Antonino Nicoletti, Grégory Franck, Stéphane Illiano

**Author notes:** Equal contribution to the manuscript (Antonino Nicoletti, Grégory Franck, Stéphane Illiano).

## Abstract

**Context:** Fever is an evolutionarily conserved and adaptive response during infections. However, prolonged fever has numerous systemic metabolic and functional side effects. In the heart, prolonged fever associated with infection is linked to fatal adverse effects, particularly involving impaired coronary circulation. Yet, the direct relationship between elevated temperature and coronary microcirculation dysfunction, remains to be fully demonstrated. In this study, we aimed to explore the specific responses of human coronary artery endothelial cells (HCAECs) to fever-range hyperthermia.

**Method:** HCAECs were cultured at either 37°C or 40°C for up to 24 hours. Transcriptomic and proteomic profiles were obtained through microarray and mass spectrometry after 6, 12, and 24 hours of exposure. Key signaling pathways, upstream regulators, and candidate mechanisms were identified and validated at the mRNA and protein levels using mechanistic approaches.

**Results:** Prolonged hyperthermia compromised HCAEC function, evidenced by cell detachment, loss of junctions, and increased permeability. HCAECs rapidly activated the unfolded protein response (UPR), including IRE1α activation and *XBP1* splicing. Additionally, autophagic flux was significantly elevated, participating in the degradation of junction proteins. Pharmacological inhibition of IRE1α reduced the autophagic burden, protected against cell detachment, and preserved junction integrity over time.

**Conclusion:** Our findings reveal that in response to fever-range hyperthermia, the IRE1α-autophagy axis regulates the survival and function of coronary endothelial cells. This mechanism could play a key role in modulating endothelial responses during infection and contribute to the pathological outcomes of fever. Furthermore, it may be relevant in local inflammatory conditions with elevated temperatures.

## Introduction

Temperature is a physical factor essential to life, governing the rates of biological and chemical reactions, particularly those catalyzed by enzymes (1). In humans, physiological temperature variations occur naturally, such as during the day-night cycle (36–37.5°C) (2) and the menstrual cycle (+0.5°C after ovulation) (3). However, febrile episodes, where body temperature rises to 38–41°C, represent a significant deviation from normal ranges, particularly during infections.

Fever is an evolutionarily conserved response that provides a protective mechanism by limiting the spread of infectious agents such as bacteria and viruses. However, prolonged fever can lead to systemic metabolic disruptions and organ dysfunction. One concerning outcome is the cardiovascular dysfunction, where elevated body temperature has been associated with various cardiovascular events (4, 5).

In the context of infection or systemic inflammation (e.g., sepsis), prolonged fever was associated with macro- and microvasculature dysfunction in the heart (6–8), which in turn compromises oxygen delivery to cardiac tissue. At the cellular level, fever-induced microcirculation impairment, characterized by endothelial hyperpermeability, disseminated intravascular coagulation (DIC), and thrombo-inflammation, has been observed in severe infections (9) (10). These observations suggest that fever could be an independent predictor of endothelial dysfunction in inflammatory diseases (11). However, the direct effects of prolonged hyperthermia on endothelial cells, particularly in the context of coronary and microvascular dysfunction, remain largely unexplored.

While several studies have investigated the cellular responses to extreme heat shock in the context of tumor hyperthermia therapy (12) and laser-induced retinal treatment (13), few have focused on febrile-range hyperthermia and its impact on cell function. Given that endothelial cells (ECs) are directly exposed to thermal heterogeneity due to their role in maintaining vascular homeostasis, understanding how they adapt or respond to such stress is crucial.

In this study, we aim to explore how human coronary artery endothelial cells (HCAECs) respond to febrile-range hyperthermia. By focusing on key molecular pathways such as the unfolded protein response (UPR) and autophagy, we seek to elucidate the mechanisms that drive endothelial cell dysfunction under these conditions and to identify potential therapeutic targets to mitigate the adverse effects of prolonged fever.

## Method

### Cell culture

Human coronary artery endothelial cells (HCAECs, #CC-2585, Lonza) were cultured in EBM-2 basal medium (#CC-3156, Lonza) supplemented with EGM-2 MV microvascular endothelium cell growth medium (#CC-3202, Lonza). Cells were maintained at 37°C in a 5% CO₂ humidified incubator and used between passages 3 and 6. To simulate febrile conditions, cells were cultured under static conditions at either 37°C or 40°C, depending on the experiment.

### Treatments

To investigate the molecular mechanisms underlying the response to hyperthermia, HCAECs were treated with the following compounds: B-I09 (0.2-20 µM, #6009, Tocris), an IRE1α inhibitor, to block XBP1 splicing and assess the role of the unfolded protein response (UPR); Chloroquine (100 µM, Sigma, C6628), was used to inhibit autophagy by blocking lysosomal function. Staurosporine (1 µM, #1285, Bio-Techne), was used as a pro-apoptotic-inducing agent (positive control).

### RFP-GFP-LC3B transfection

*100,000* HCAECs were grown until 70µl confluency in µ-Slide 8 Well high Glass Bottom (Ibidi), and transfected with 10µl of Premo autophagy tandem sensor RFP-GFP-LC3B mix (Molecular Probes, P36239) diluted in 300 µl of culture medium per well. Transfected cell were grown overnight in normal conditions and then cultured at 37°C or 40°C for 3 or 24h. After nuclei stain with Hoechst (1:1000), and mounting with 200µl ProLong Gold antifade reagent, both RFP and GFP positive puncta were quickly visualized by using an AxioObserver microscope (Zeiss, x63) and images were processed and quantified with Fiji (ImageJ) software for quantification.

### RNA Isolation and Microarray Analysis

For transcriptomic analysis, HCAECs were seeded at a density of 5 × 10⁵ cells per well in 6-well plates and cultured in complete medium for 6, 12, or 24 hours at 37°C or 40°C. Total RNA was extracted using the miRNEasy mini kit (#217004, Qiagen) according to the manufacturer’s instructions. RNA quality was assessed using the Agilent 2100 Bioanalyzer, and microarray analysis was performed using the SurePrint G3 Human Gene Expression v3 8×60K Microarray Kit (Agilent). Differential gene expression was determined by comparing the 40°C and 37°C conditions, and significant changes were defined as a fold change of ≥±2 with a p-value < 0.05, using Partek Flow (Illumina). Pathway analysis was performed using (Ingenuity Pathway Analysis, Qiagen).

### qRT-PCR

Total RNA was reverse-transcribed using the SuperScript VILO cDNA Synthesis Kit (#11754050, Invitrogen). qRT-PCR was carried out with TaqMan Universal PCR Master Mix (#4369016, Invitrogen) on a Stratagene Mx3000P detection system (Agilent). Primer sets used in our study are listed in the **Supplemental Table 1**. Relative gene expression was calculated using the 2^−ΔΔCt^ method.

### Western Blot

Cells were lysed in ice-cold cell extraction buffer (#FNN0011, Invitrogen) supplemented with protease and phosphatase inhibitors (#P8340 and #P0044 Sigma). Protein concentration was determined using the Pierce BCA Protein Assay Kit (#23225, Thermo Fisher). Proteins (1–10 µg per sample) were separated by SDS-PAGE and transferred to nitrocellulose or PVDF membranes for subsequent immunoblotting. Primary antibodies were used against sXBP1, LC3B, p62, VE-Cadherin and phospho-VE-Cadherin (**Supplementary Table 2**). Detection was performed using HRP-conjugated appropriate secondary antibodies and chemiluminescence substrate (GE Healthcare).

### Immunofluorescence

After treatments, cells were fixed and permeabilized with Cytofix/Cytoperm solution (#554714, Becton Dickinson). Immunolabeling was performed using primary antibodies against LC3B, LAMP2, VE-Cadherin, and cleaved caspase-3 (**Supplementary Table 2**). After incubation with secondary fluorescent antibodies (Thermo Fisher), nuclei were stained with Hoechst (1:1000), and slides were mounted with ProLong Gold antifade reagent. Images were captured using an AxioObserver microscope (Zeiss) and processed with Fiji (ImageJ) software for quantification.

### Electron microscopy

Cells were fixed 1h in a mix of 1% glutaraldehyde, 2,5% paraformaldehyde in PBS, then washed 3 x 5 minutes in PBS. Samples were then post-fixed during 1h in reduce Osmium tetroxide 1% Potassium Ferrocyanide 1,5%, then washed 3 x 5 minutes in water. Dehydration was performed using graded series of ethanol in water for 10 minutes 30%, 50%, 70%, 80%, 90% (x2), 100% (x3). Resin infiltration was performed by incubating 1h in an Agar low viscosity resin (Agar Scientific Ltd) and ethanol mix (1:1), followed by overnight incubation in pure resin at 4°C. The resin was then changed and the samples further incubated during 3 hours prior to flat inclusion in gelatin capsules and overnight polymerization at 60°C. 70 nm sections were obtained using an EM-UC6 ultramicrotome (Leica), post-stained in 4 % aqueous uranyl acetate and lead citrate, and observed at 120 kV with a Tecnai12 transmission electron microscope (ThermoFisher Scientific) equipped with a 4K×4K Oneview camera (Gatan).

### Statistical Analysis

Statistical analysis was performed using GraphPad Prism 9.0. Data are presented as mean ± standard error of the mean (SEM), and comparisons between two groups were made using the Mann-Whitney U test. A p-value < 0.05 was considered statistically significant. N represents the number of different experiments performed on different days.

## Results

### Prolonged febrile range hyperthermia impairs HCAEC survival adherence and function

HCAECs cultured at 40°C exhibited significant structural and functional alterations compared to cells maintained at 37°C. As early as 6 hours, transmission electron microscopy revealed the formation of large lysosomal structures (**Fig. 1A**). After 12 hours, cells began to lose their junctions, and after 24 hours, there was a marked detachment of cells, along with the formation of apoptotic bodies and of microparticles (**Fig. 1A**). Quantification of VE-Cadherin staining confirmed progressive junctional disruption over time at 40°C (**Fig. 1B**). A significant increase in apoptosis was observed in flow cytometry, with Annexin V-positive cells rising by 30% at 12 hours and by 20% at 24 hours in the 40°C condition (**Fig. 1C**). Staurosporine was used as a positive control in these experiments and the impact of temperature was 55 and 40 % of the maximal value obtained with staurosporine at 12 and 24h, respectively. Additionally, cleaved-caspase 3 staining indicated increased apoptotic signaling in cells lacking continuous junctions (**Fig. 1D**). Supernatant analysis confirmed a steady rise in both detached cells and apoptotic bodies or microparticles after 24 hours of culture at 40°C (**Fig. 1E**).

**Figure 1:**
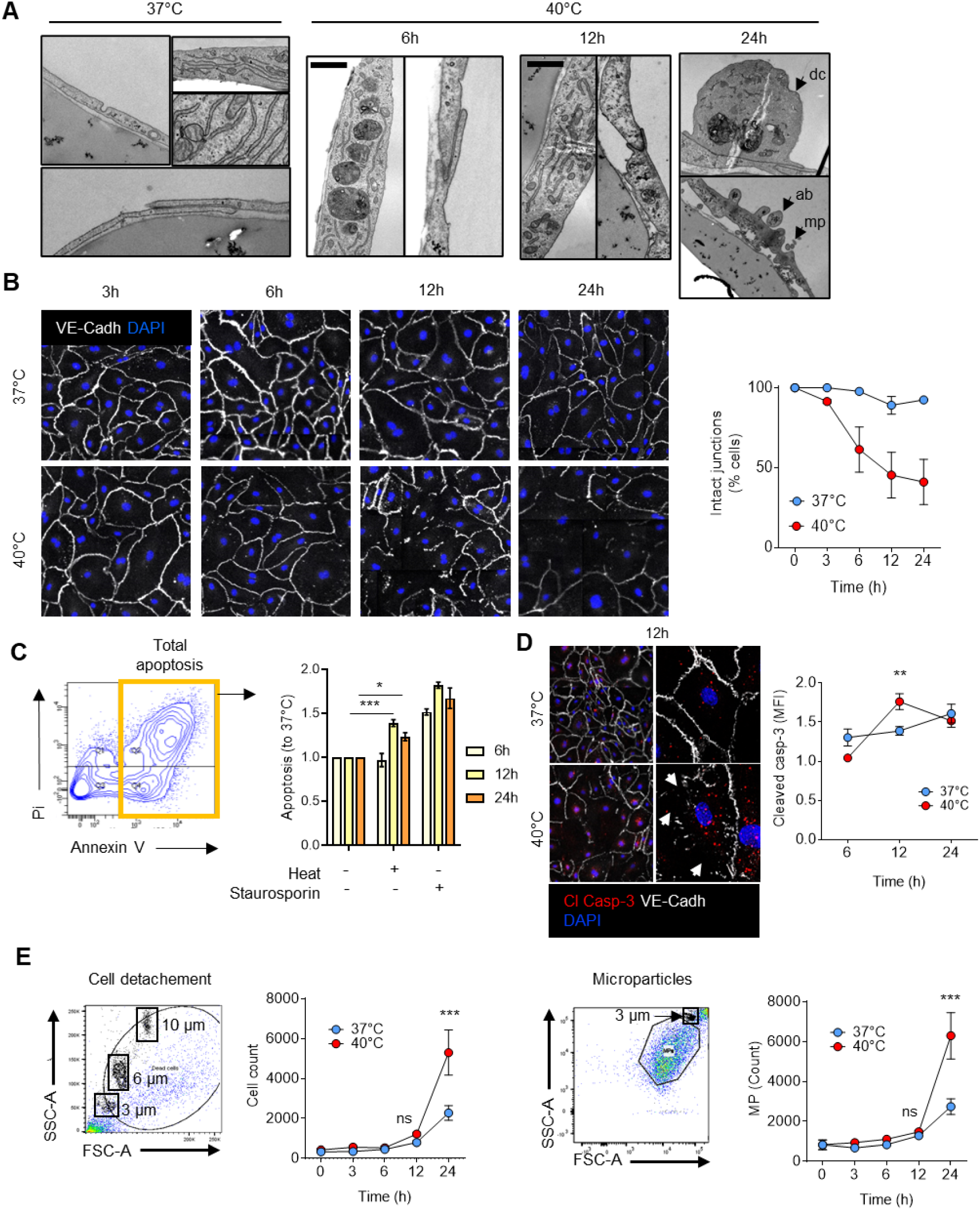
Febrile range hyperthermia impairs EC survival, adherence and function *in vitro*. **A.** Ultrastructural analysis (TEM, down) of HCAECs subjected to hyperthermia for 6, 12 or 24h. **B.** Cell-cell junction shown by VE-Cadherin staining over time in HCAECs subjected or not to FRH (left) and semi-quantitated (right). **C.** Apoptosis levels evaluated by flow cytometry using Pi Annexin V staining in HCAECs subjected or not to FRH for 6, 12 and 24h. **D.** VE-cadherin and cleaved caspase-3 immunostaining in HCAECs subjected to FRH for 12h. Corresponding semi quantitative analysis for cleaved caspase-3 signal (MFI) in HCAECs subjected or not to FRH. Arrowheads show cell junction perturbations. **E.** Number of detached cells (left) and microparticles (right) in the media of HCAECs subjected to FRH for 3, 6, 12 and 24h, evaluated by flow cytometry. Each observation is made of n=3 independent experiments. * p<0.05; **p<0.01; ***p<0.001.

### Transcriptomic and proteomic analyses reveals molecular mechanisms underlying the response to febrile-range hyperthermia

To dissect the molecular basis of these phenotypic changes, we conducted transcriptomic and proteomic profiling of HCAECs exposed to 40°C. Microarray analysis revealed that 243, 607, and 532 mRNA probes were downregulated at 6, 12, and 24 hours, respectively, while 378, 267, and 286 probes were upregulated (**Fig. 2A and 2B**). Principal component analysis (PCA) indicated a clear segregation of gene expression patterns between the 37°C and 40°C conditions, with significant clustering based on temperature (**Fig. 2C**). The most significantly upregulated pathways included the unfolded protein response (UPR) and endoplasmic reticulum (ER) stress, with IRE1α and XBP1 identified as key early upstream regulators (**Fig. 2D, 2E, and Supplementary Fig. 1**). The effect of FRH was also investigated at the proteomic level, using an ion trap mass spectrometry analysis (Orbitrap). Proteomic analysis indicated an enrichment of proteins involved in selective autophagy, adherens junctions, and ER-related processes (**Fig. 2F and 2G**). Differentially expressed proteins (DEPs) identified through an ANOVA analysis are summarized in a volcano plot comparing cells cultured at 37°C and 40°C (**Fig. 2F**, Fold change >1.5). The identified DEPs (10 downregulated, 24 upregulated) allowed a biological process enrichment analysis of most significant modules of the protein-protein interaction (PPI) network (CluegGO, Cytoscape) and showed that early response to fever-range hyperthermia upon proteomic profiling involved selective autophagy (52%), along with adherens junctions (4%), interconversion of nucleotide di-and triphosphate (8%), ADP metabolic process (8%), eukaryotic 48S preinitiation complex (12%), and snRNP assembly (16%) (**Fig. 2G**).

**Figure 2:**
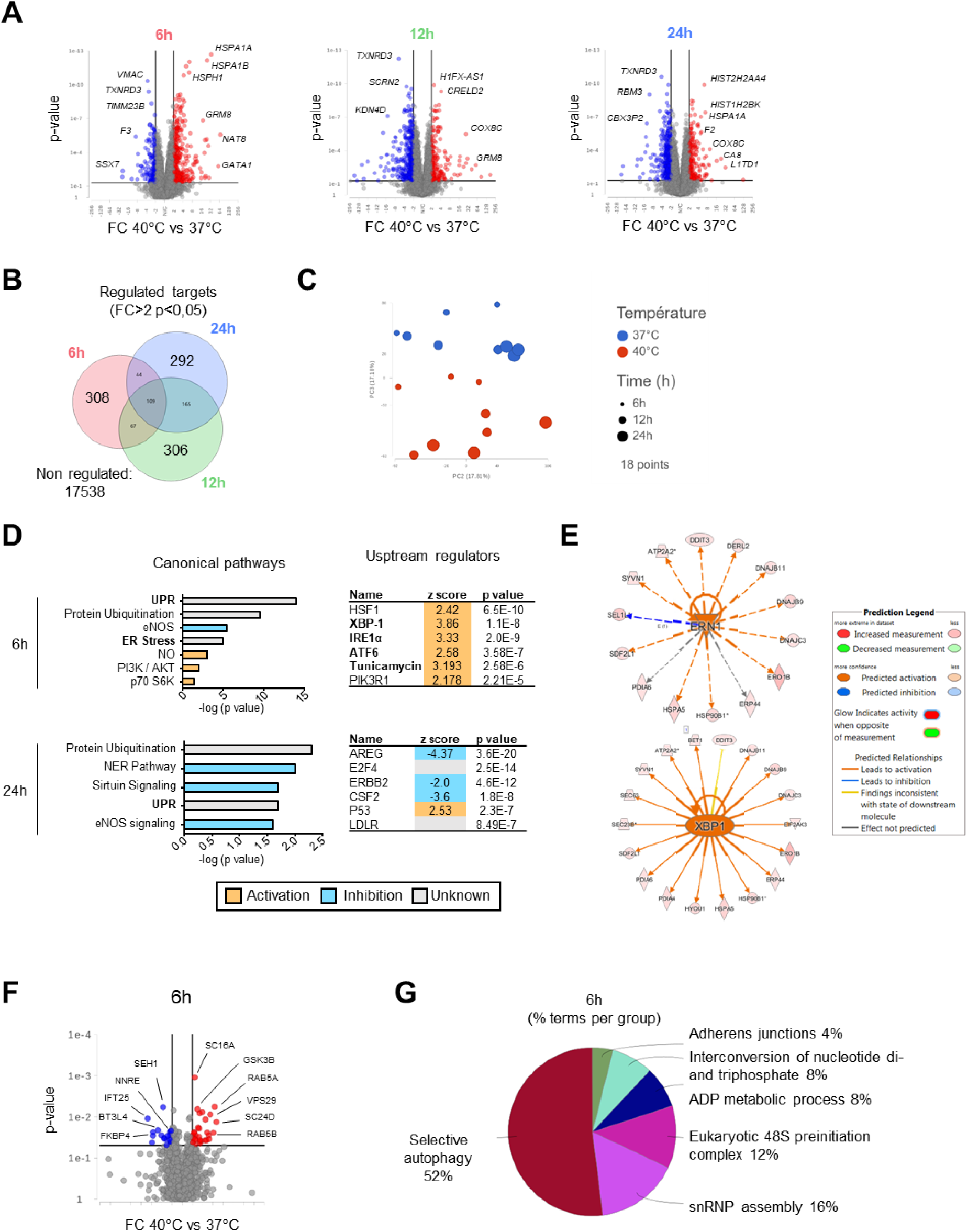
Unbiased transcriptomic and proteomic analysis unravels molecular mechanisms involved in EC response to febrile range hyperthermia. **A.** Volcano plots illustrating the differentially expressed genes (DEGs) over time (40°C *vs* 37°C). The most significant DEGs are highlighted. **B.** Venn diagram showing the relationship between regulated targets over time (40°C *vs* 37°C). **C.** Principal component analysis color-coded for temperature. Dot size represents different time points. Predicted canonical pathways activated at 6h and 24h (**D**) and most significant upstream regulators identified (**E**). **F.** Volcano plot illustrating the differentially expressed proteins (DEPs) at 6h (40°C *vs* 37°C). The most significant DEPs are highlighted. **G** GO/pathway terms enrichment for the given DEPs.

### ER stress and IRE1α are activated in response to fever-range hyperthermia

Electron microscopy showed signs of ER stress, with notable swelling of the endoplasmic reticulum in cells cultured at 40°C (**Fig. 3A**). qRT-PCR analysis revealed that the expression of ER stress-associated genes such as *GRP78* and *CHOP* was transiently upregulated between 2 and 6 hours (**Fig. 3B**). IRE1α activation was assessed by quantifying both spliced (s) and unspliced (u) *XBP1* mRNA. s*XBP1* levels increased significantly at 1 and 3 hours in the 40°C condition, while u*XBP1* levels remained unchanged (**Fig. 3C**). The transient increase in XBP1 protein was confirmed and quantified in western Blot. **Fig. 3D-E**). Treatment with the IRE1α inhibitor B-I09 effectively blocked XBP1 splicing in a dose-dependent manner (**Fig. 3F**). Among several targets of the UPR, the expression of *DNAJB9*, a transcript coding a member of the DNAJ (Hsp40) family with various proteostatic functions, was significantly decreased after treatment by B-I09 (**Supplementary Fig. 2A**). Of note, hyperthermia also activated the PERK branch of UPR, as the phosphorylation of EiF2α was found higher in cells cultured at 40°C (**Supplementary Fig. 2B**).

**Figure 3:**
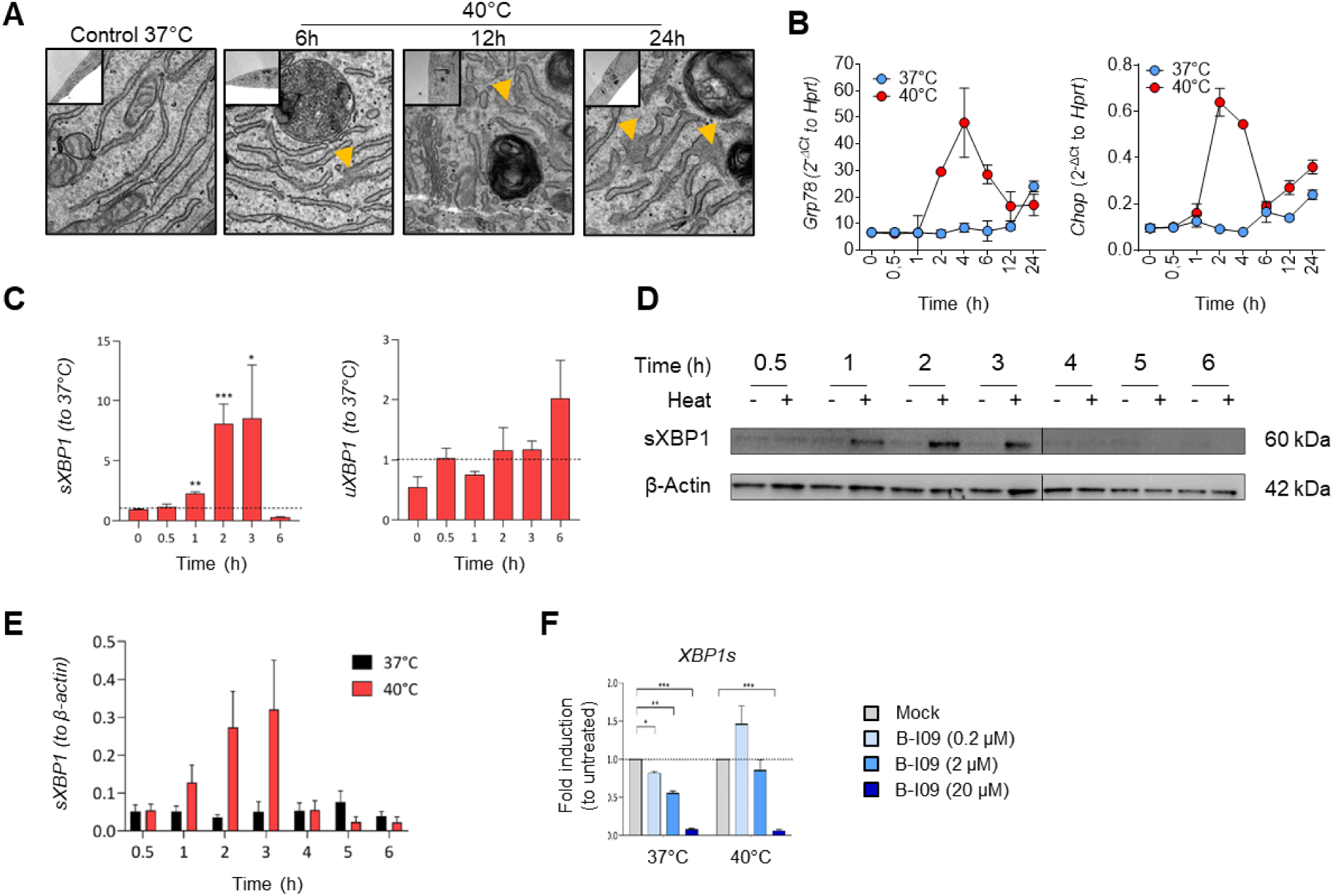
ER stress response and XBP1 splicing in HCAECs subjected to FRH. **A.** Ultrastructural analysis of HCAEC by TEM showing endoplasmic reticulum swelling over time. These pictures are representative of 10 pictures / conditions, in 3 independent experiments. **B.** Expression kinetic of *GRP78* et *CHOP* mRNAs over 24h during FRH. **C.** Expression kinetic of *spliced (s)XBP1* and *unspliced (u)XBP1* mRNA over 6 hours during FRH as compared to 37°C. **D.** Western blot representative of XBP1s protein kinetic expression in HCAECs exposed to moderate hyperthermia (40°C vs 37°C). β-actin was used as a loading control. This blot is representative of 3 independent experiments. Quantitative analysis of the kinetic of XBP1s protein expression in HCAECs exposed to 37°C or 40°C (N=3 experiments). **F.** RT-qPCR showing the spliced XBP1 (XBP1s) or unspliced XBP1 (XBP1u) mRNA abundance in HCAEC cultured at 37°C or 40°C for 2h with or without B-I09 (20µM) (N=3 experiments). *P<0.05, **P<0.01, ***P<0.001

### Hyperthermia increases autophagy flux in ECs

Hyperthermia-induced autophagy was investigated using LC3B immunostaining and Western blot analysis. HCAECs cultured at 40°C exhibited a modest but significant increase in LC3B expression over time (**Fig. 4A**, n=3). LC3B increase was further amplified by chloroquine treatment, which blocks lysosomal degradation, indicating enhanced autophagic flux. p62 levels, a marker of autophagic degradation, were also elevated at 3, 6, and 12 hours in the 40°C condition, confirming the increased autophagic activity (**Fig. 4B**). Immunofluorescence staining revealed co-localization of LC3B^+^ autophagosomes with LAMP2^+^ lysosomes, suggesting active autophagosome-lysosome fusion during hyperthermia (**Fig. 4C**, n=10 per condition). RFP-GFP-LC3B transfection experiments confirmed transient autophagy induction, with a significant increase in autolysosome formation at 3 hours, which normalized by 24 hours (**Fig. 4D**). HCAEC cultured at 40°C harbored abnormal number of amphisome-like structures carrying high number of autophagosomes, lysosomes and possibly endosomes (**Fig. 4E**). In parallel, cells had increased vesicle trafficking along stages of FRH, suggesting frequent endocytosis and a membrane protein turnover (**Fig. 4F**). Immunostaining revealed colocalization of LC3B^+^ autophagosomes with VE-Cadherin when cells were cultured at 40°C, suggesting that internalization vesicles are taken over by autophagy (**Fig. 4G**). Altogether, these data show that temperature dysregulates autophagy, increasing its flux in an extent that could be detrimental for the cell function and survival.

**Figure 4:**
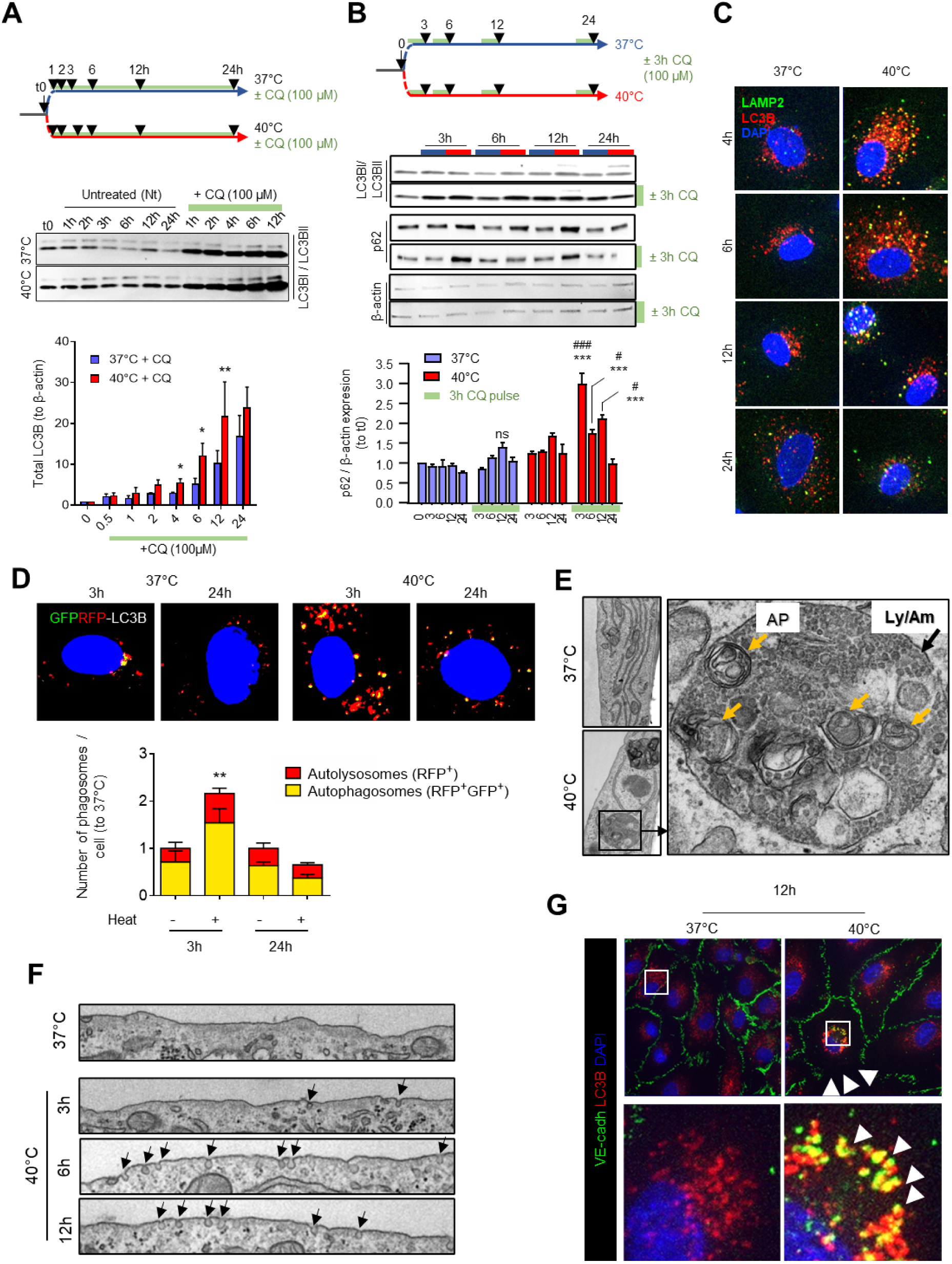
Heat transiently increases autophagy flux in HCAECs. **A.** Experimental design and LC3B levels by western blot to assess autophagy flux in HCAEC in response or not to FRH, in presence or absence of 100µM chloroquine (CQ). Semi-quantitative measurement is shown on the bottom. **B.** Experimental design and LC3B or p62 levels by western blot to assess autophagic flux within a defined time window by sequential CQ addition. Semi-quantitative measurement is shown on the right. **C.** LAMP2 / LC3B co-immunolabelling in HCAEC subjected to FRH over time (n=10 per condition. Images representative of 3 different experiments). **D.** Fluorescence detection of GFP-RFP-LC3B transfected in HCAEC subjected or not to FRH during 3 or 24h, with autolysosomes in red (RFP+) and autophagosomes in yellow (GFP+RFP+). Semi-quantitative measurement is shown on the right. **E.** Cytosol ultrastructure in HCAEC subjected to FRH for 12h. **F.** Plasma membrane ultrastructure in HCAECs subjected to FRH over time. Arrows indicate endosomal formation. **G.** VE-Cadherin / LC3B co-immunolabelling in HCAEC subjected or not to FRH for 12h. Arrowheads point at colocalization of autophagosomes with VE-Cadherin. * p<0.05, ** p<0.01, *** p<0.001, 40°C *vs* 37°C in CQ-treated cells. # p<0.05, ### p<0.001, CQ-treated cells *vs* untreated cells in group 40°C. Student t-test.

### IRE1α inhibition alleviates hyperthermia-induced autophagy and preserves junction integrity

To evaluate the role of IRE1α in hyperthermia-induced dysfunction, we treated HCAECs with the IRE1α inhibitor B-I09 (0.2-20µM). Inhibition of IRE1α effectively prevented cell detachment and reduced microparticle and apoptotic body production after 24 hours at 40°C (**Fig. 5A-B**). Interestingly, B-I09 did not reduce heat-mediated apoptosis, suggesting that cell detachment is not directly mediated by apoptotic mechanisms (**Fig. 5C**). Additionally, B-I09 treatment normalized autophagy flux, as evidenced by reduced LC3B levels and decreased autophagosome formation (**Fig. 5D and 5E**). Western blot analysis confirmed that B-I09 treatment restored VE-Cadherin levels, and immunofluorescence showed improved junction integrity compared to untreated cells during hyperthermia (**Fig. 5F-G**). Collectively, these data demonstrate that febrile-range hyperthermia impairs endothelial cell function, leading to cell detachment, junctional disruption, and increased autophagy. Inhibition of IRE1α by B-I09 mitigates these effects by preventing cell detachment, reducing the autophagic burden, and restoring the integrity of cellular junctions, suggesting a critical role of the IRE1α-autophagy axis in maintaining endothelial cell stability under thermal stress (**Fig. 5H**).

**Figure 5:**
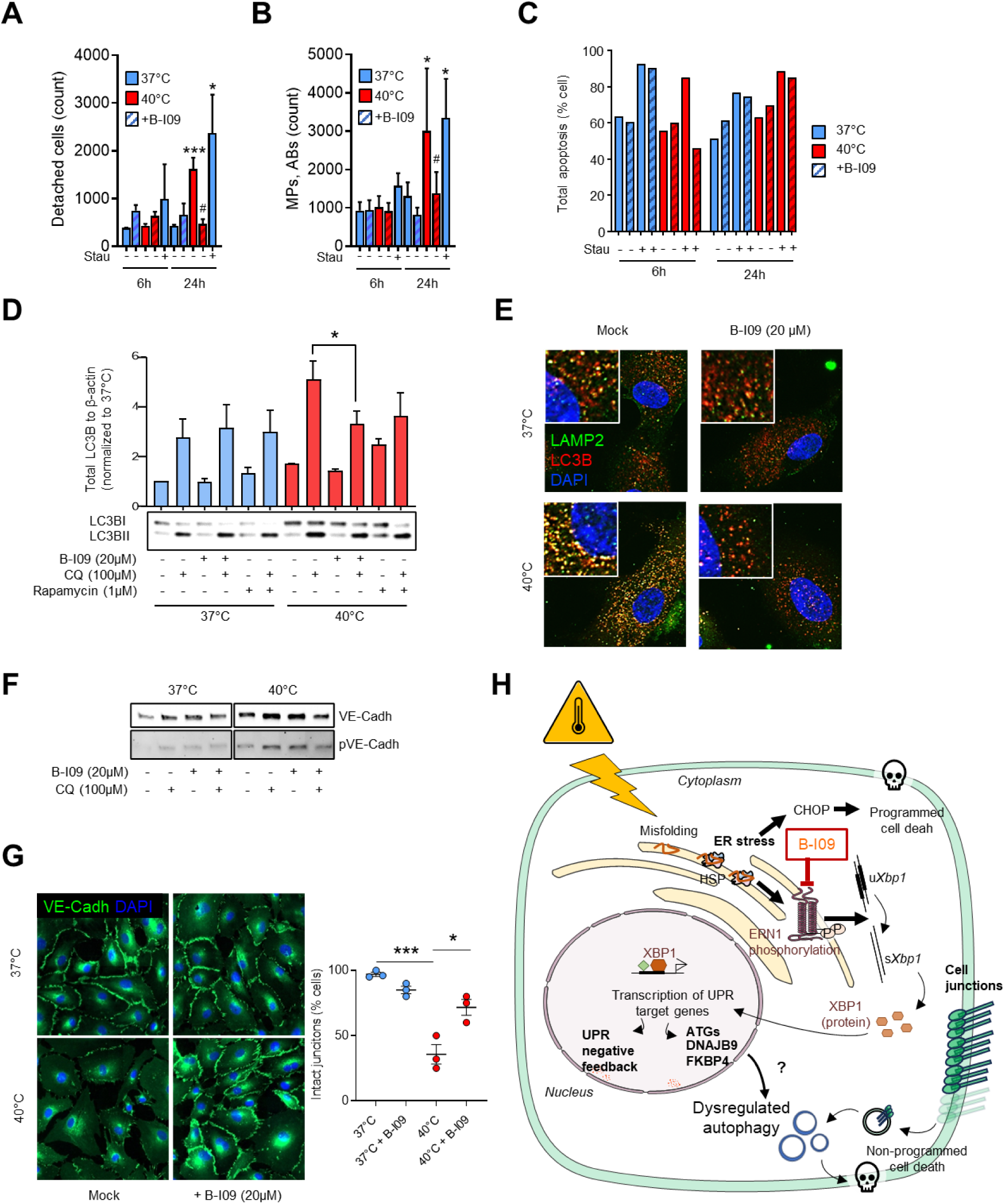
IRE1 inhibition by B-I09 alleviates HCAEC detachment and junction alteration in response to FRH. Effect of B-I09 on cell detachment (**A**) and microparticle production (**B**) and apoptosis (**C**) by HCAECs in response to FRH. Stau: staurosporine. *p<0.05, ***p<0.001 (40°C vs 37°C), # p<0.05 (B-I09 *vs* mock). Effect of B-I09 in HCAECs subjected to FRH for 6h on LC3B levels (western blot, **D**) or autophagosome / autolysosome presentation (LC3B/LAMP2 immunofluorescence, **E**). **F.** Levels of total VE-Cadherin assessed by western blot in presence of chloroquine (CQ) or B-I09 in HCAECs in response to FRH for 6h. **G.** Cell junction quality assessed by VE-Cadherin immunolabeling in HCAEC subjected or not to FRH. Corresponding semi quantitative analysis showing intact junction percentage on the right. **H.** Summary diagram of the study. *p<0.05, ***p<0.001 between the indicated groups, by student unpaired t-test.

## Discussion

Our study provides novel insights into how febrile-range hyperthermia (FRH) triggers a coordinated cellular stress response in coronary endothelial cells, with a central role for the unfolded protein response (UPR) and autophagy. While fever is recognized as a beneficial evolutionary response during infections, prolonged febrile episodes can lead to significant cardiovascular complications, including endothelial dysfunction, microvascular hyperpermeability, and thrombo-inflammation (10),(11). Our findings identify the IRE1α-XBP1 axis as a key regulator of endothelial cell survival under febrile stress, connecting ER stress to autophagic pathways that have been proposed to control cellular junction integrity and function (14).

Previous studies have largely focused on the response of endothelial cells to extreme heat shock (13), leaving a gap in understanding the effects of more physiological temperature elevations such as FRH. Our work bridges this gap by demonstrating that endothelial cells exhibit early ER stress, as evidenced by XBP1 splicing and IRE1α activation, within hours of exposure to 40°C. Notably, we show that this stress response is closely linked to an autophagic burst, a finding that complements previous studies on the role of autophagy in endothelial stress responses (15). Our data align with work by Margariti *et al.* (15), who also identified the IRE1α-XBP1 pathway as an inducer of autophagy in endothelial cells, but extend their findings by connecting this response to cell detachment and junction disruption during hyperthermia.

These results have direct implications for clinical conditions associated with prolonged fever, such as sepsis or severe viral infections, where endothelial dysfunction plays a critical role in disease progression (16–20). In these settings, fever-driven endothelial cell damage could exacerbate microvascular impairment, contributing to organ failure and adverse outcomes. Our finding that IRE1α inhibition by B-I09 preserves endothelial cell junctions and prevents detachment offers a potential therapeutic avenue for protecting the vasculature during febrile episodes. Pharmacological targeting of IRE1α could be explored as a strategy to mitigate endothelial dysfunction and its complications in diseases characterized by systemic or localized fever, such as sepsis, Kawasaki disease, or even certain autoimmune conditions (e.g. lupus erythematosus).

Moreover, the preservation of VE-Cadherin junctions in our study underscores the potential to maintain vascular integrity under thermal stress, which could be particularly relevant for patients with pre-existing endothelial vulnerabilities, such as those with cardiovascular disease or diabetes.

### Limitations of the study

This study primarily relies on *in vitro* models of hyperthermia in cultured endothelial cells, which, while valuable, may not fully recapitulate the complexity of in vivo microvascular dynamics. Future research should include in vivo models to validate these findings, particularly in the context of infection-induced fever and its effects on systemic vascular health. *In vitro*, some experiments using flow conditions could be used to validate these finding, since shear stress plays an important role in EC biology and survival. Additionally, while B-I09 proved effective in our experiments, the broader physiological consequences of IRE1α inhibition remain to be explored, as this pathway plays a role in various cellular processes beyond autophagy and ER stress management.

### Perspectives

Given the multifaceted role of the UPR in endothelial cells, further research should explore the involvement of other UPR branches, such as PERK and ATF6, in the cellular response to hyperthermia. It will be important to elucidate how these pathways interact with autophagy and whether they contribute to endothelial dysfunction in the same manner as IRE1α. Moreover, future studies should aim to identify the long-term consequences of modulating UPR pathways, particularly in chronic febrile conditions or in diseases where persistent low-grade inflammation is coupled with endothelial stress.

Another promising avenue of research involves testing the effects of IRE1α inhibition in vivo, both in animal models of febrile infection coupled with microvascular dysfunction and to extend it to more complex models with vascular inflammation.

## Conclusion

This study reveals that prolonged febrile-range hyperthermia impairs endothelial cell function by triggering a complex response involving the activation of the unfolded protein response (UPR) and a significant increase in autophagic activity. Our results demonstrate that the IRE1α-XBP1 axis plays a central role in mediating these effects, linking ER stress to cellular junction disruption and detachment. Importantly, pharmacological inhibition of IRE1α not only mitigates these detrimental effects but also preserves cell junction integrity and reduces the autophagic burden, highlighting a potential therapeutic target for maintaining endothelial function under febrile conditions. These findings have significant implications for clinical conditions characterized by prolonged fever, such as sepsis and severe viral infections (21), where endothelial dysfunction is a critical factor in disease progression. Future research should focus on validating these findings in vivo and exploring the long-term consequences of IRE1α inhibition, particularly in the context of inflammatory diseases. Additionally, the interplay between other branches of the UPR and autophagy remains to be fully elucidated, offering further opportunities to understand and modulate endothelial responses to thermal stress.

---

**Supplementary Figure 1.**
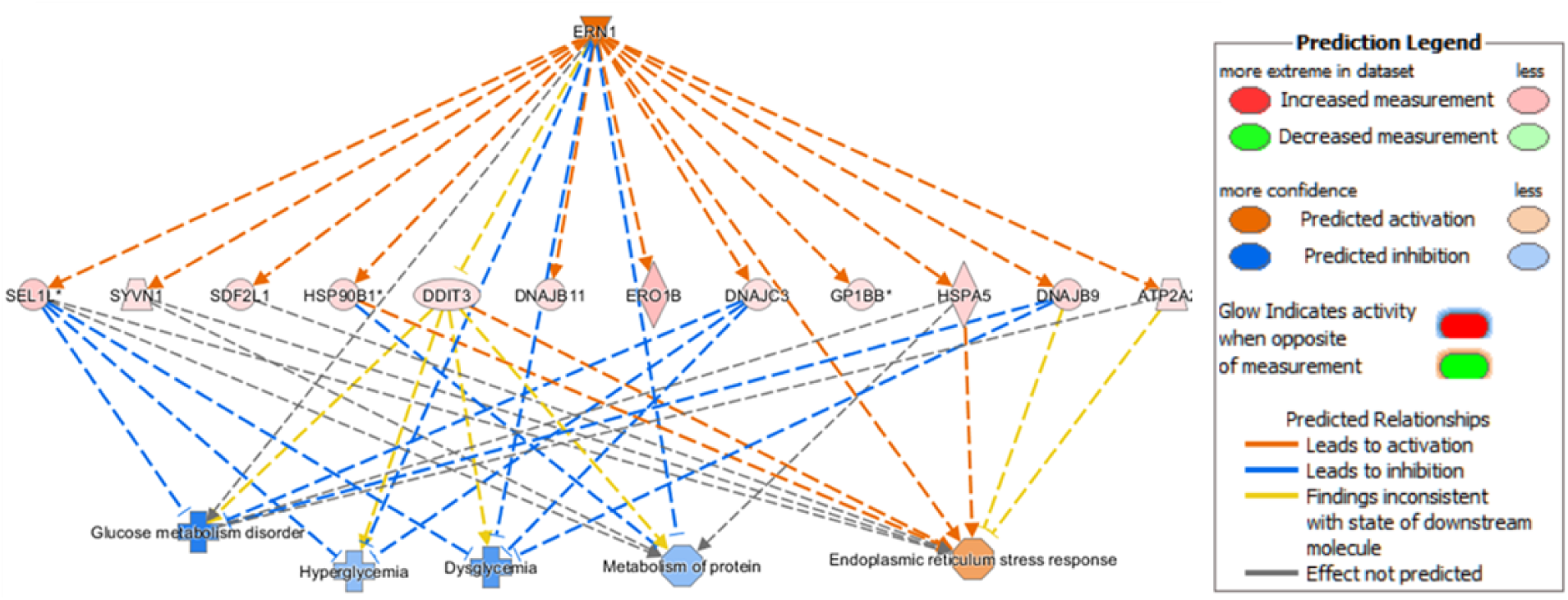
Top1 putative upstream regulator ERN1 (IRE1α) and its known targets.

**Supplementary Figure 2.**
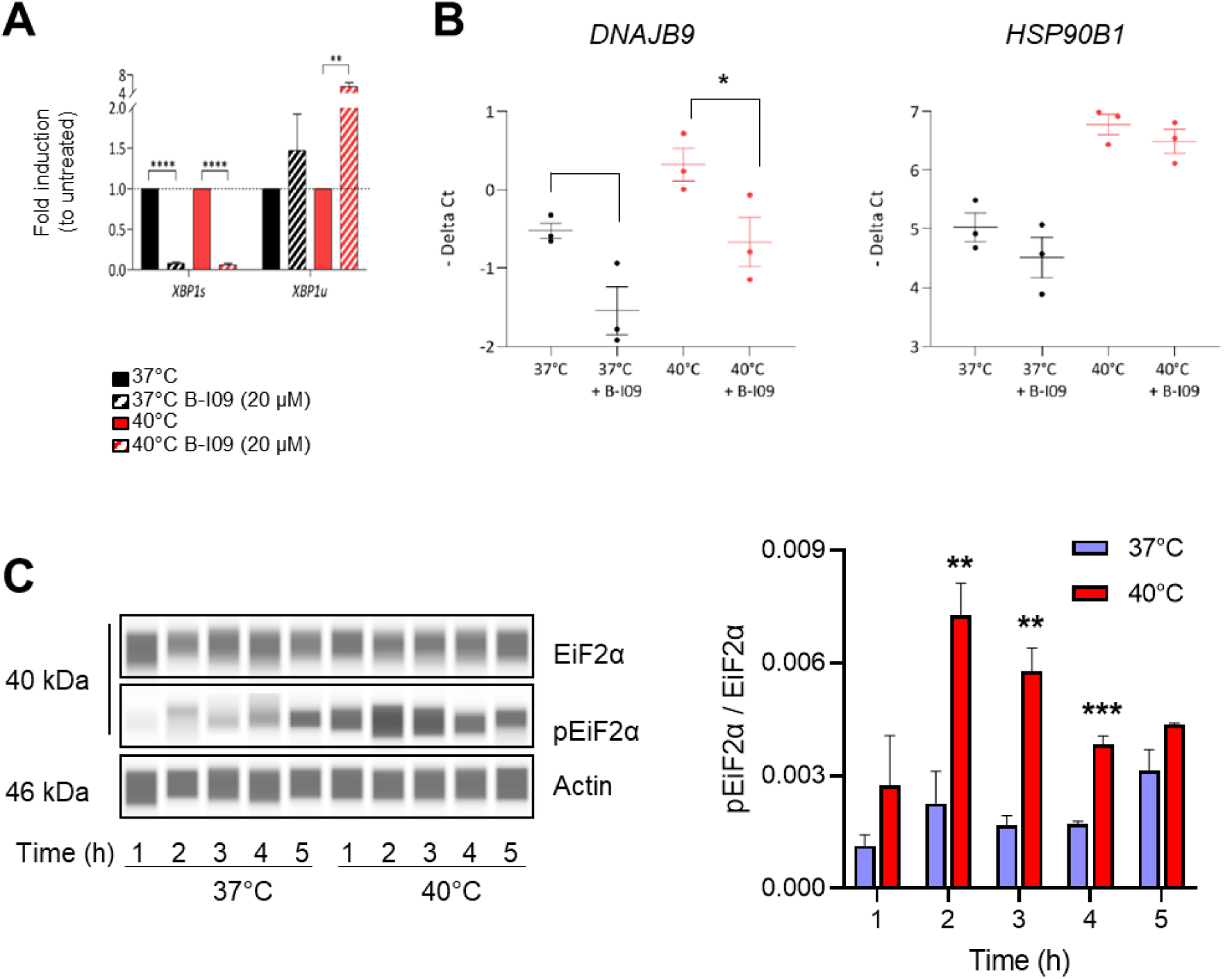
Effect of FRH on the unfolded protein response in HCAECs. **A.** Effect of B-I09 (20 µM) on sXBP1 and uXBP1 mRNA levels after 2h at 37°C or 40°C. **B.** Effect of B-I09 on the mRNA levels of *DNAJB9* and *HSP90B1*. **C.** Level of phosphorylation of EiF2α in HCAECs subjected or not to FRH, assessed by Wes Simple Western System, and quantified (right).

**Supplemental Table 1.**
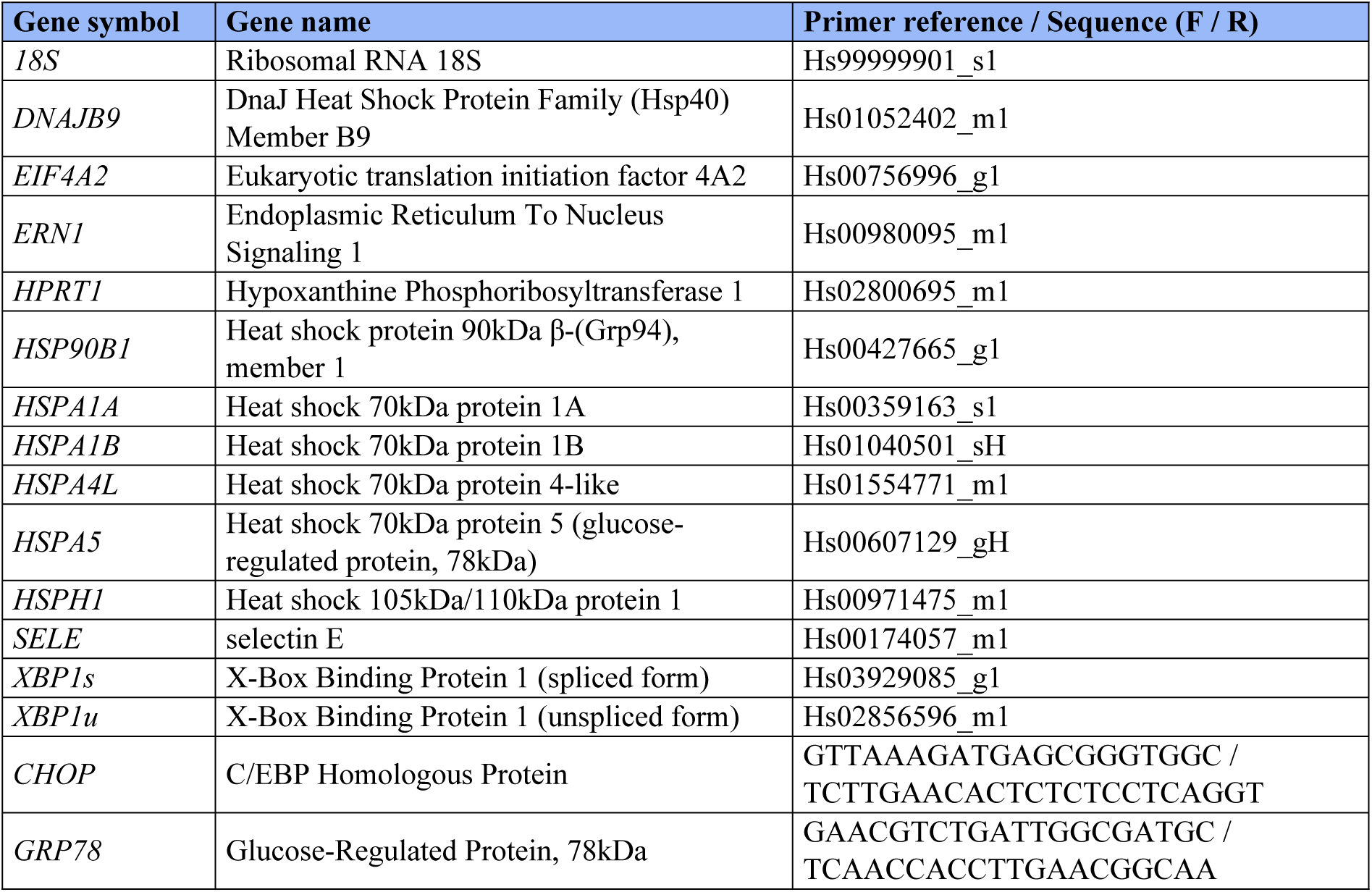

**Supplemental Table 2.**
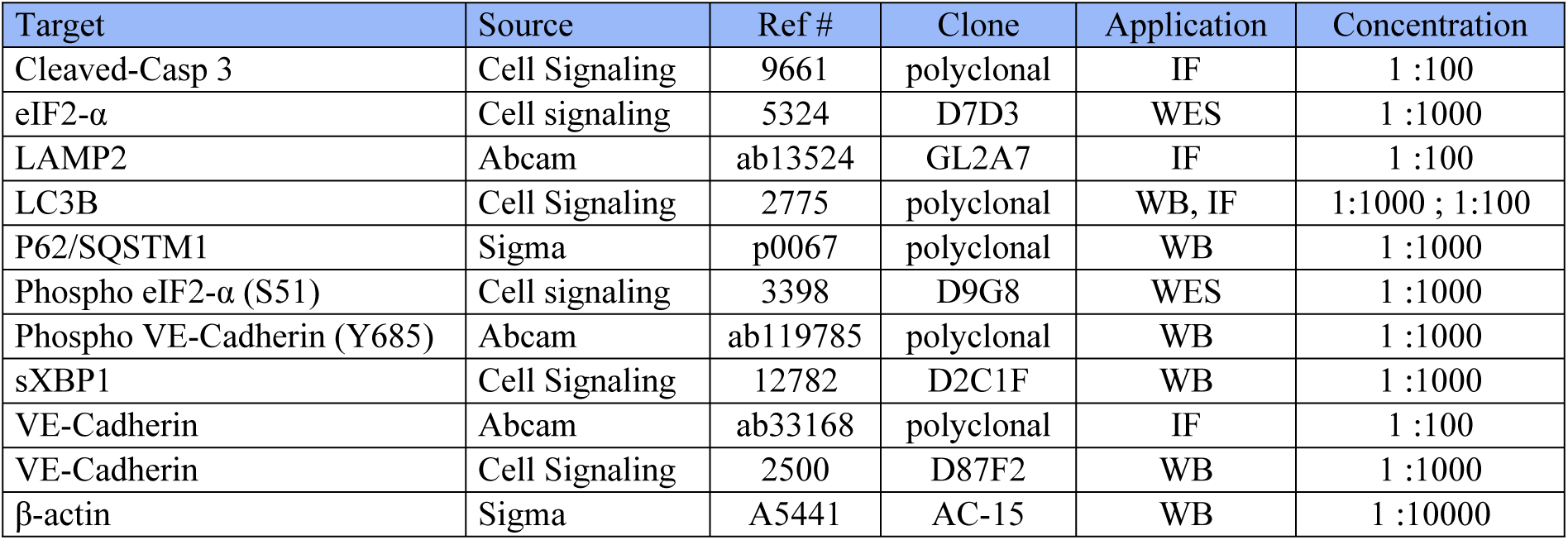
Antibodies used for western blot and immunofluorescence.

## Acknowledgment

We acknowledge Rémi Le Borgne and the ImagoSeine core facility of the Institut Jacques Monod, member of the France BioImaging infrastructure (ANR-10-INBS-04) and GIS-IBiSA, for their assistance with cell processing and the transmission electron microscopy observations. We thank Guillaume Chevreux and the Proteomics Core Facility (Institut Jacques Monod UMR 7592) for their assistance with proteomic analysis.

For the purpose of Open Access, a CC-BY public copyright license has been applied by the authors to the present document distributed under a Creative Commons Attribution | 4.0 International license (https://creativecommons. org/licenses/by/4.0/).

## References

1. Peterson ME, Daniel RM, Danson MJ, Eisenthal R. The dependence of enzyme activity on temperature: determination and validation of parameters. Biochem J. 2007;402(2):331–7.

2. Van Someren EJ. Mechanisms and functions of coupling between sleep and temperature rhythms. Prog Brain Res. 2006;153:309–24.

3. Tur E. Physiology of the skin--differences between women and men. Clin Dermatol. 1997;15(1):5–16.

4. Manger B, Schett G. Aortic dissection and fever: cause or consequence. Z Rheumatol. 2017;76(6):550–1.

5. Auala T, Zavale BG, Mbakwem AC, Mocumbi AO. Acute Rheumatic Fever and Rheumatic Heart Disease: Highlighting the Role of Group A Streptococcus in the Global Burden of Cardiovascular Disease. Pathogens. 2022;11(5).

6. Walter EJ, Hanna-Jumma S, Carraretto M, Forni L. The pathophysiological basis and consequences of fever. Crit Care. 2016;20(1):200.

7. Kakihana Y, Ito T, Nakahara M, Yamaguchi K, Yasuda T. Sepsis-induced myocardial dysfunction: pathophysiology and management. J Intensive Care. 2016;4:22.

8. Maneta E, Aivalioti E, Tual-Chalot S, Emini Veseli B, Gatsiou A, Stamatelopoulos K, et al. Endothelial dysfunction and immunothrombosis in sepsis. Front Immunol. 2023;14:1144229.

9. Li XK, Yang ZD, Du J, Xing B, Cui N, Zhang PH, et al. Endothelial activation and dysfunction in severe fever with thrombocytopenia syndrome. PLoS Negl Trop Dis. 2017;11(8):e0005746.

10. de Sousa FTG, Warnes CM, Manuli ER, Ng A, D’Elia Zanella L, Ho YL, et al. Yellow fever disease severity and endothelial dysfunction are associated with elevated serum levels of viral NS1 protein and syndecan-1. medRxiv. 2023.

11. Mori Y, Katayama H, Kishi K, Ozaki N, Shimizu T, Tamai H. Persistent high fever for more than 10 days during acute phase is a risk factor for endothelial dysfunction in children with a history of Kawasaki disease. J Cardiol. 2016;68(1):71–5.

12. Pelz J, Mollwitz M, Stremmel C, Goehl J, Dimmler A, Hohenberger W, et al. The impact of surgery and mild hyperthermia on tumor response and angioneogenesis of malignant melanoma in a rat perfusion model. BMC Cancer. 2004;4:53.

13. Du S, Zhang Q, Zhang S, Wang L, Lian J. Heat shock protein 70 expression induced by diode laser irradiation on choroid-retinal endothelial cells in vitro. Mol Vis. 2012;18:2380–7.

14. Reglero-Real N, Perez-Gutierrez L, Yoshimura A, Rolas L, Garrido-Mesa J, Barkaway A, et al. Autophagy modulates endothelial junctions to restrain neutrophil diapedesis during inflammation. Immunity. 2021;54(9):1989–2004 e9.

15. Margariti A, Li H, Chen T, Martin D, Vizcay-Barrena G, Alam S, et al. XBP1 mRNA splicing triggers an autophagic response in endothelial cells through BECLIN-1 transcriptional activation. J Biol Chem. 2013;288(2):859–72.

16. Caliskan M, Gullu H, Yilmaz S, Erdogan D, Unler GK, Ciftci O, et al. Impaired coronary microvascular function in familial Mediterranean fever. Atherosclerosis. 2007;195(2):e161–7.

17. Li XK, Zhang SF, Xu W, Xing B, Lu QB, Zhang PH, et al. Vascular endothelial injury in severe fever with thrombocytopenia syndrome caused by the novel bunyavirus. Virology. 2018;520:11–20.

18. Portier I, Campbell RA, Denorme F. Mechanisms of immunothrombosis in COVID-19. Curr Opin Hematol. 2021;28(6):445–53.

19. Iba T, Levy JH, Levi M. Viral-Induced Inflammatory Coagulation Disorders: Preparing for Another Epidemic. Thromb Haemost. 2022;122(1):8–19.

20. Mussbacher M, Salzmann M, Brostjan C, Hoesel B, Schoergenhofer C, Datler H, et al. Cell Type-Specific Roles of NF-kappaB Linking Inflammation and Thrombosis. Front Immunol. 2019;10:85.

21. Aljadah M, Khan N, Beyer AM, Chen Y, Blanker A, Widlansky ME. Clinical Implications of COVID-19-Related Endothelial Dysfunction. JACC Adv. 2024;3(8):101070.

